# Visualizing Long-Term Single-Molecule Dynamics *in vivo* by Stochastic Protein Labeling

**DOI:** 10.1101/116186

**Authors:** Hui Liu, Peng Dong, Maria S. Ioannou, Li Li, Jamien Shea, H. Amalia Pasolli, Jonathan Grimm, Pat Rivlin, Luke D. Lavis, Minoru Koyama, Zhe Liu

## Abstract

Our ability to unambiguously image and track individual molecules in live cells is limited by packing of multiple copies of labeled molecules within the resolution limit. Here we devise a universal genetic strategy to precisely control copy number of fluorescently labeled molecules in a cell. This system has a dynamic titration range of >10,000 fold, enabling sparse labeling of proteins expressed at different abundance levels. Combined with photostable labels, this system extends the duration of automated single-molecule tracking by 2 orders of magnitude. We demonstrate long-term imaging of synaptic vesicle dynamics in cultured neurons as well as in intact *zebrafish*. We found axon initial segment utilizes a ‘waterfall’ mechanism gating synaptic vesicle transport polarity by promoting anterograde transport processivity. Long-time observation also reveals that transcription factor hops between clustered binding sites in spatially-restricted sub-nuclear regions, suggesting that topological structures in the nucleus shape local gene activities by a sequestering mechanism. This strategy thus greatly expands the spatiotemporal length scales of live-cell single-molecule measurements, enabling new experiments to quantitatively understand complex control of molecular dynamics *in vivo*.

## INTRODUCTION

Life is orchestrated by animate molecular processes in living cells. Decoding the dynamic behavior of single molecules enables us to investigate biology at this fundamental level that is often obscured in ensemble experiments. Sparse labeling is critical for live-cell single-molecule imaging^1, 2^, because high copy numbers (10^3^ ~ 10^7^) of protein molecules co-occupy the small volume in a cell (10 ~ 20 μm in diameter). Thus, multiple protein molecules are packed within the light diffraction limit (200 ~ 300 nm). For example, in a small area (10 μm × 10 μm) comparable to the size of a mammalian cell nucleus, single-molecule images begin to overlap with each other at low concentrations (> 10^2^ ~ 10^3^ copies), preventing precise localization (**Supplementary Fig. 1a, b**). Accurate single-molecule tracking requires labeling at much lower densities (< 10^2^ copies) to link single-molecule positions across multiple frames with high fidelity (**Supplementary Fig. 1b**). Currently, the most reliable live-cell single-molecule imaging method is by stochastic switching of photoactivatable fluorescent proteins or dyes (sptPALM or dSTORM)^2^-^9^. The stochastic photoactivation separates appearances of individual fluorophores temporally, allowing single-molecule imaging in densely labeled samples. However, under these conditions, only one fluorophore per molecule can be utilized for imaging. Thus, high laser powers (> 10^3^ W/cm^2^) potentially damaging to live specimens are applied to generate sufficient photon counts for localization^10^. As a result, fluorescent molecules are quickly bleached and single-molecule dynamics can only be probed on relatively short time scales. Recently, a new method (MINFLUX) is developed to minimize the photon flux for accurate localization, enabling long-term tracking of single photoactivatable proteins in live bacteria. However, highly-focused scanning beams limit the field of view (50 ~ 100 nm)^11^. Fast imaging of a larger object such as a human cell and simultaneous tracking of multiple molecules are still challenging^12^.

Here we reason that if we can devise a genetic strategy to precisely control copy number of fluorescently labeled proteins in a cell, this would allow us to sparsely label any target protein with bright fluorescent tags (*e.g.* 3XGFP, 3XHaloTag, GFP11^13^, sfCherry11^13^, SunTag^14^ and ArrayG^15^) for long-term single-molecule imaging and tracking. One unique advantage of this method is that single-molecule imaging across a large field of view can be realized by conventional microscopy without photoactivation. Currently, a common strategy to control labeling density is by using promoters with variable strengths to adjust mRNA transcript levels. However, there are no guidelines to engineer the strength of a promoter as promoter strengths are cell-type dependent^16^ and weak promoters are associated with high gene expression noise^17, 18^. These disadvantages make it challenging to fine tune protein copy numbers by transcriptional control.

Here, we first demonstrate a general method to precisely control copy number of fluorescently labeled proteins by translational readthrough (RT). With this stochastic labeling strategy, we could sparsely label densely-packed cellular organelles and proteins with bright fluorescent labels for long-term single-molecule imaging. Real-time imaging in neurons reveals a critical role of axon initial segment in enhancing synaptic vesicle anterograde movement. Long-term observation of transcription factor (Sox2) binding dynamics show that topological structures in the nucleus kinetically facilitate TF molecules to translocate among clustered binding sites, confining local target search and gene regulatory activities. Our studies thus provide the first evidence supporting a dynamic sequestering mechanism of topological domains in the nucleus of live cells.

## RESULTS

### Stochastic Protein Labeling with a Translational Readthrough Strategy

During protein translation, ribosome sometimes mis-incorporates an amino acid at the stop codon and continues to translate the downstream mRNA sequence. The RT mechanism is conserved from bacteria to mammals^19^-^21^. Interestingly, the probability that a ribosome translates beyond the stop codon (the RT efficiency) is largely dictated by the sequence of the stop codon and its immediate flanking regions^20, 21^. Simulation experiments suggest that, compared to using weak promoters, sparse labeling by a RT strategy would benefit from averaging effect at the translational level and give rise to much less labeling heterogeneity in single cells (**Supplementary Fig. 2**). To test this approach, we performed live-cell PALM^22^ experiments using cells expressing Histone H2B and mEOS4b separated by different RT sequences (**Fig. 1a, b**). Although mRNAs were expressed at similar levels (**Supplementary Fig. 3c**), we observed dramatic reductions of mEOS4b labeling densities upon the RT sequence insertion (**Fig. 1b, c, Supplementary Fig. 3b** and **Supplementary Movie 1**). We found that each RT sequence has a distinct labeling probability (RT1; ~5%, RT2; ~1.5%, RT3; ~0.4%) (**Fig. 1c**), similar to the RT efficiency determined for each sequence by luciferase assays^21^.

**Figure 1:**
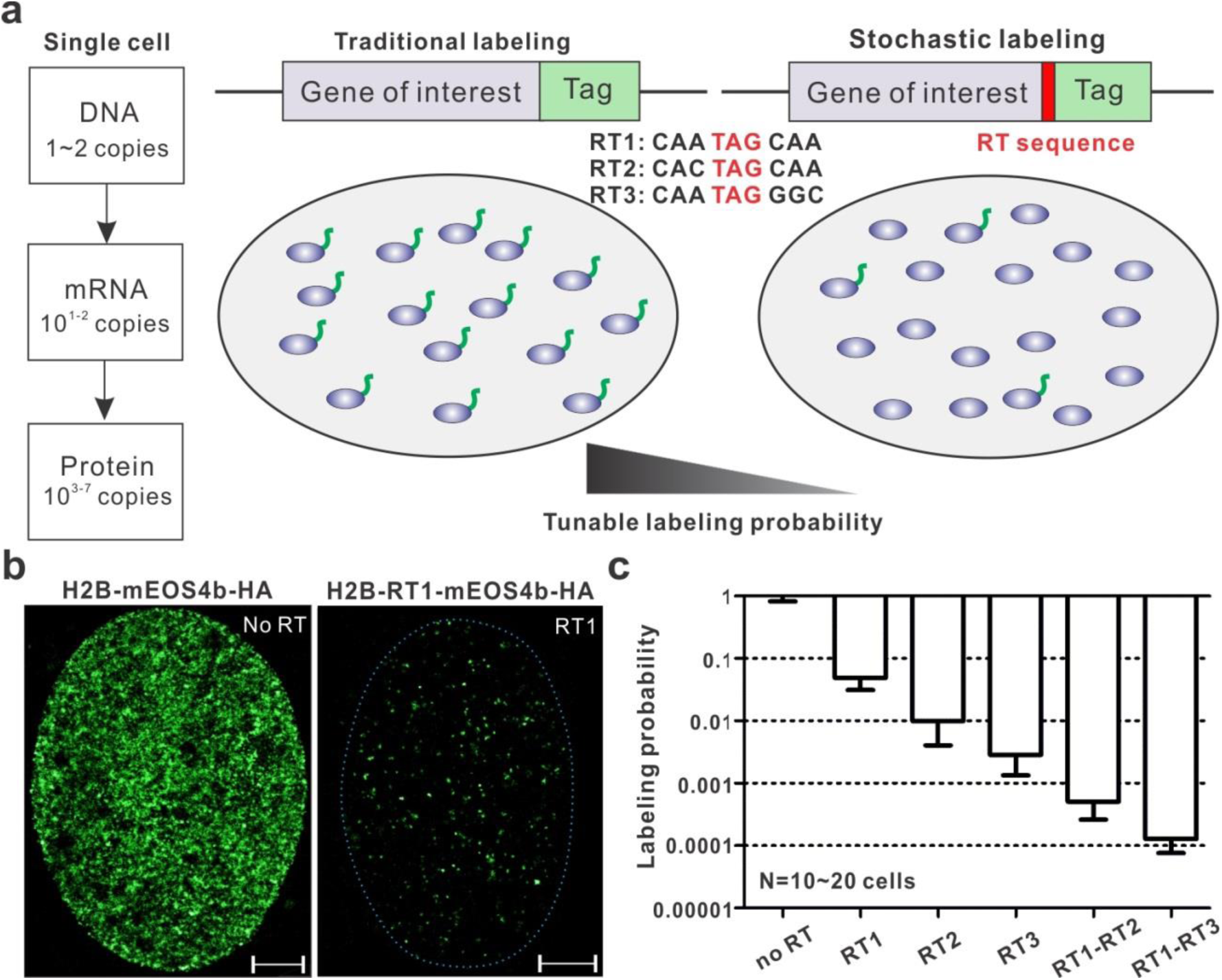
Sparse Labeling with Translational Read-Through. (**a**) Schematics for precise label density control by translational read-through Left: After transcription and translation, high copy numbers of functional protein molecules (10^3^ ~ 10^7^) are typically generated in the cell. Middle: In the traditional labeling strategy, all molecules are labeled and high labeling densities make it challenging to observe single-molecule dynamics (**Supplementary Fig. 1**) Right: By placing a short RT sequence between the target protein and the tag, we can accurately control the probability of protein fusion and sparsely label a faction of molecules for singlemolecule imaging. (**b**) Representative single-cell PALM localization density map for H2B-mEOS4b in U2OS cells without (Left) and with (Right: RT1) read-through (**Supplementary Movie 1**). In average, 328 localizations per μm^2^ (no RT) and 17.2 localizations per μm^2^ (RT1) were detected within 4500 frames. Scale bar: 2 μm. (**c**) Labeling probabilities for different RT sequences have a large dynamic range (from ~5% down to ~0.001%). Specially, for the probability calculation, H2B-mEOS4b localization density in each RT condition were normalized to the average density in no RT control. Error bars represent standard deviation (SD).

Importantly, these probabilities are consistent across 4 different cell types that we tested (**Fig. 1c** and **Supplementary Fig. 3d**), suggesting that the RT system functions with similar efficiencies in different cell-types. We also discovered that RT sequences work synergistically in tandem, giving rise to even lower labeling probabilities (RT1-RT2; ~0.06%, RT1-RT3; ~0.01%) (**Fig. 1c**). Therefore, the RT system can titrate labeling density linearly on the logarithmic scale with a large dynamic range (~10,000 fold). If necessary, this range can be further extended by concatenation of RT sequences. In contrast, the internal ribosome entry site (IRES) only reduced the labeling density to ~40% (**Supplementary Fig. 3a**), not suitable for sparse labeling of most proteins in a cell. It is important to note that past studies identified a repertoire of short sequences with RT efficiencies ranging from 0.1% up to ~10%^23^. These sequences can also be adapted to make fine adjustments to the labeling probability. Based on these data, we propose that the highly tunable RT system can be used to stochastically label protein in live cells for single-molecule imaging (**Fig. 1a**).

### Long-term Observation of Synaptic Vesicle Moving Dynamics

We first sought to apply this stochastic labeling strategy to an important question in neuroscience - how neuronal trafficking polarity is dynamically established. Specifically, synaptic vesicle precursors (SVPs) packaged at the soma must be specifically targeted to presynaptic regions for release and recycle as synaptic vesicles (SVs)^24^. Airyscan super-resolution imaging (**Fig. 2b** and **Supplementary Fig. 4a**) and electron microscopy (EM) studies^25, 26^ reveal that high copy numbers of SVPs/SVs are densely packed below the diffraction limit in mammalian neurons, particularly in boutons and presynaptic regions. Consistent with these results, when we labeled a classical synaptic vesicle marker – Synaptophysin (Syp) with traditional strategies (*e.g.* CMV*d3* and SunTag v1), individual SVPs/SVs cannot be distinguished from each other (**Fig. 2a**, **b** and **Supplementary Fig. 5**). Next, we implemented the RT strategy to sparsely label Syp with the SunTag (**Fig. 2a** and **Supplementary Fig. 4b**) - a fluorescence signal amplification system capable of directing 24 copies of fluorescence protein to the target^14^. After optimizing the RT sequence, we can directly observe movements of bright, sparsely-spaced SVPs/SVs in different compartments of the cell (**Fig. 2b** and **Supplementary Movie 2**). The advantages for this strategy are *1)* there are ~30 copies of Syp in each SVP/SV^27, 28^ and on average only one copy of Syp is labeled (**Fig. 2a**).

**Figure 2:**
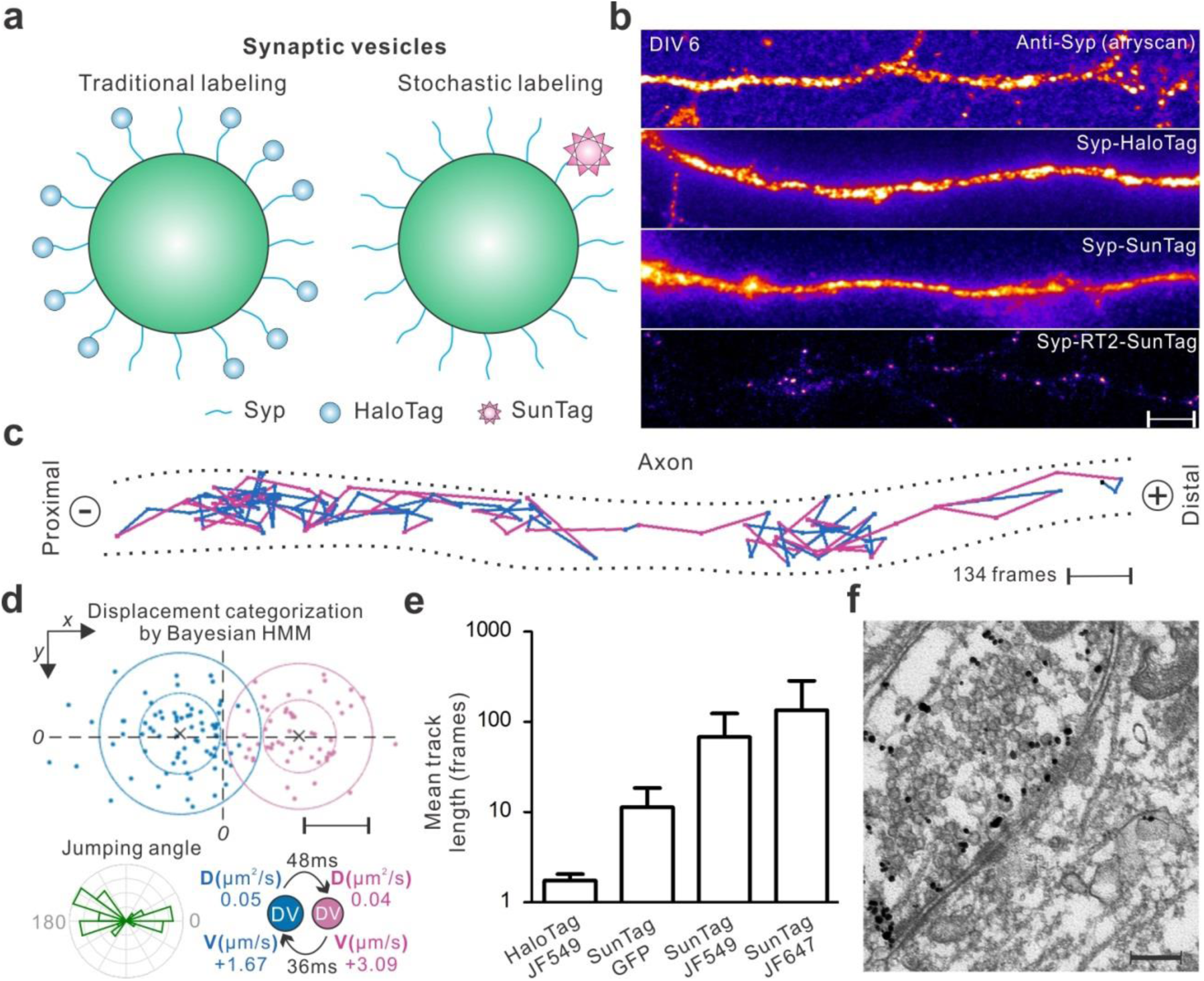
Automated long-term tracking of individual synaptic vesicle precursors in live neurons. (**a**) SVP/SV labeling either by tagging Syp with the HaloTag (Left) or sparsely with the SunTag via RT sparse labeling (Right). (**b**) High packing densities prevent imaging single SVP/SV particles with traditional labeling methods in DIV 6 cultured hippocampal neurons (From top to bottom, Anti-Syp staining (Airyscan); HaloTag fusion (total internal reflection fluorescence - TIRF); SunTag without RT sparse labeling (TIRF)). When sparsely labelled with the SunTag (lower panel, TIRF) using RT2, individual and sparsely-spaced SVPs/SVs (scFv-HaloTag-JF646) can be observed and imaged. DIV is short for Days In Vitro. Scale bar: 5 μm (**c**) One representative trajectory tracking SVP movements in axon: steps are color-coded according to transport modes classified by the HMM-Bayes algorithm shown in (**d**). The ‘+’ and ‘-’ denote anterograde and retrograde transport directions, respectively. Scale bar: 200 nm. See **Supplementary Movie 3** for raw imaging data. (**d**) Displacements between frames for the trajectory shown in (**c**) can be classified into two main categories with one transport component pointing to the ‘+’ direction and the other to the ‘-’ direction. Bottom left, Rose histogram for jumping angles for trajectory shown in (**c**). Bottom right, transport velocities - V, diffusion coefficients - D and mean lifetimes for the two states defined in the upper panel. The transport process is modeled as directed motion with diffusion, according to <*r*^2^>= 4*D*Δ*t* + (*V*Δ*t*)^2^^46^ (See *Methods* for calculation details). Scale bar: 100 nm (**e**) Track length comparison for different labeling strategies. Data in lane 1 were acquired using the dSTORM strategy with high laser powers (>1 kW/cm^2^). Data in Lane 2 ~ 4 were acquired with relatively low laser powers (~50 W/cm^2^) with sparse SunTag labeling. Error bars represent SD. (**f**) Immuno-EM images showing SunTag labeled SVPs/SVs (RT1) were ts in DIV argeted to presynaptic region12 hippocampal neurons. Scale bar: 200 nm

Thus, we expect that the large size (> 1.4 Md) of the SunTag would not significantly affect the SVP/SV function. Indeed, both immunofluorescence staining and high-resolution EM experiments demonstrated that SVPs/SVs labeled with the SunTag were correctly targeted to presynaptic regions (**Fig. 2f** and **Supplementary Fig. 6**). *2)* Due to sparse labeling, the movements of single SVPs/SVs covering large distances in the cell can be unambiguously imaged and tracked with high accuracy (**Fig. 2c** and **Supplementary Movie 3, 7**). *3)* Because of the signal amplification from the SunTag, ~20 fold less laser powers (~50 mW/cm^2^) can be used compared to the sptPALM or dSTORM method for fast single-molecule imaging (50 Hz). Thus, imaging induced photo-toxicity is greatly reduced without trading off imaging speed. *4)* By incorporating HaloTag and bright Janelia Fluor (JF) dyes^5^ into the SunTag system, average trajectory lengths showed ~100 fold improvement over the traditional single HaloTag strategy (**Fig. 2e** and **Supplementary Movie 2**). To further validate the applicability of this method, we sparsely labeled SVPs/SVs in live *zebrafish* using the Syp-SunTag RT system (**Supplementary Fig. 7a**). We can observe dynamic transport of single SVPs along neurites *in vivo* (**Supplementary Fig. 7b, c** and **Supplementary Movie 5, 6**).

### Enhanced Anterograde Transport of SVP in Axon Initial Segment

It is worth noting that SVP/SV diffusion dynamics has been imaged before with traditional GFP labeling^29, 30^ or extracellular loading of antibody-dye conjugates^31, 32^. However, because of dense labeling, these methods do not allow long-term, fast tracking of SVPs/SVs, which is essential for quantifying SVP/SV transport dynamics. Since reliable classification of transitions between active transport and random diffusion requires continuous analysis of displacements in long trajectories^33, 34^, we applied a recently developed HMM-Bayes algorithm to analyze our SVP/SV tracking data^33^. This program effectively separates SVP/SV diffusion, anterograde and retrograde transport processes (Fig. 2c, d, **3b** and **Supplementary Fig. 7c**) and provides detailed kinetic information, such as diffusion coefficients, transport velocities and lifetimes for each state. Here for the first time, long-term dynamics of individual SVPs/SVs can be imaged and quantitatively analyzed in live mammalian neurons with high spatiotemporal resolution.

**Figure 3:**
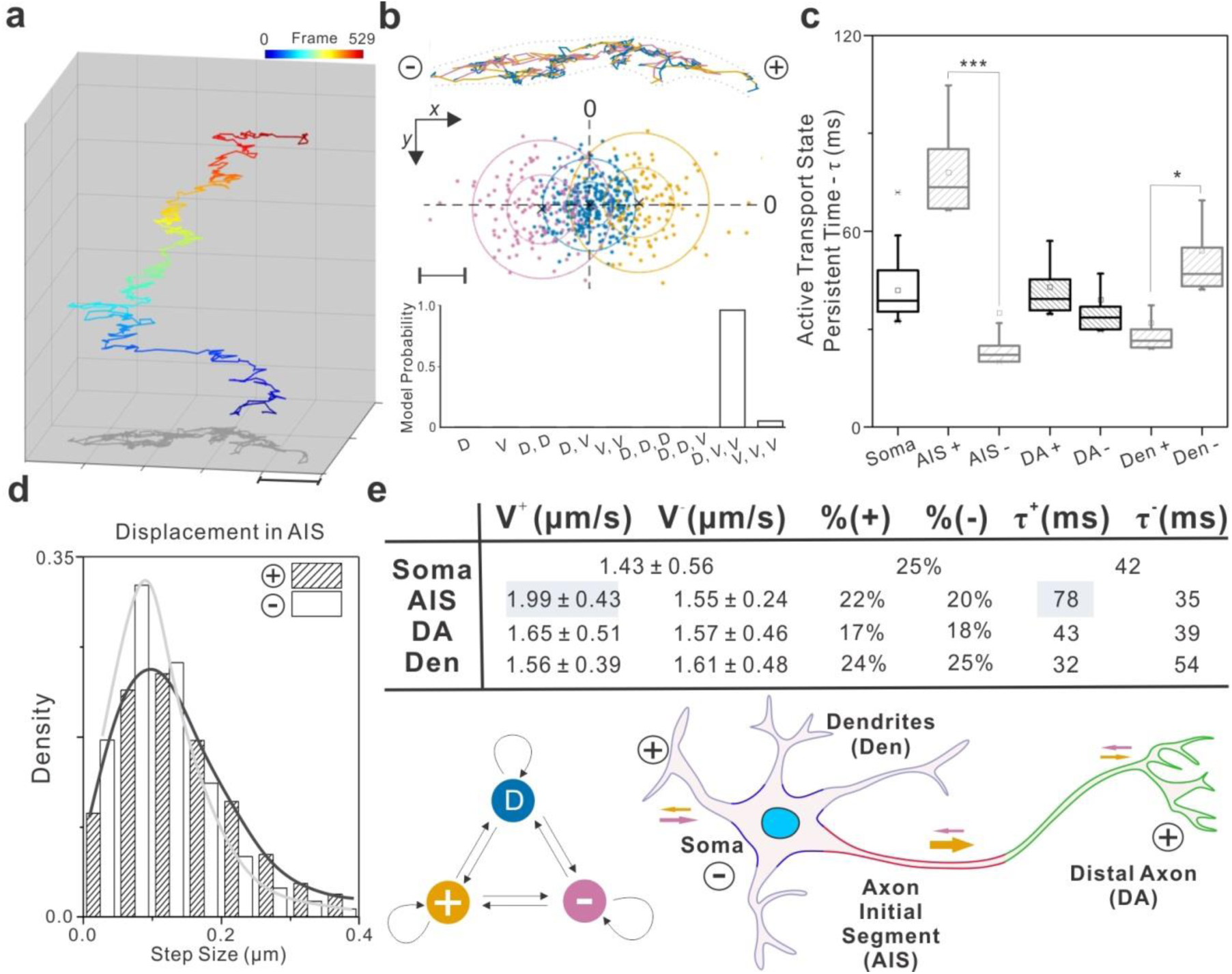
Axon initial segment gates synaptic vesicle precursor transport polarity. (**a**) A representative SVP trajectory consists of 529 frames showing dynamic transitions between diffusion and active transport states. See **Supplementary Movie 7** for the raw imaging data. The frame number is color-coded as indicated by the color bar above. Scale bar: 1 μm (**b**) Top, individual steps in the trajectory showed in (**a**) are color-coded by the three hidden states classified in the middle panel. Middle, displacements between frames for the trajectory shown in (**a**) can be classified by the HMM-Bayes algorithm into one diffusion state (blue) and two transport states with opposing directions - anterograde (+, yellow) and retrograde (-, pink). Bottom, the probabilities of competing models for correctly describing the hidden states in the trajectory. D stands for diffusion and V for active transport. Scale bar: 100 nm (**c**) The persistent lifetime for each active transport state in the indicated compartment. The abbreviations (AIS, DA and Den) are the same as defined in (**e**), ‘+’ and ‘-’ denote anterograde and retrograde transport respectively (Number of cells analyzed > 10, Number of trajectories analyzed > 300). In the box chart, top and bottom error bars represent 95% and 5% percentile, respectively; box represents the range from 25% to 75% percentile; center line represents the median and the small square represents mean value. “AIS +” versus “AIS -”: *p*-value < 4.5e-4, t =5.43, df = 578; “Den +” versus “Den -”: *p*-value = 0.032, t = -2.14, df = 463. ***, *p*-value < 0.001; *, *p*-value < 0.05 (**d**) The step size distribution for anterograde (+) and retrograde (-) transport processes in the AIS. (**e**) Top, statistic summary for anterograde (+) and retrograde (-) transport processes in different compartments of the neuron. V, transport velocity; %, time fraction; τ, state lifetime; the velocities were shown in mean ± SD. “AIS V^+^” versus “AIS V^−^”: *p*-value < 1.4e-5, t = 14.69, df = 578. Bottom left, hidden markov model diagram describing the dynamic behavior of SVP in the cell. Bottom right, schematics explaining the abbreviations for different functional compartments of the neuron and the directions of active transport. AIS, DA and Den stand for axon initial segment, distal axon and dendrites, respectively. The size of arrows indicates the degree of cargo transport processivity. All tracks from different compartments were taken from the same datasets to exclude the bias from stage or sample motion.

Next, we divided each neuron into four distinct functional compartments (Dendrites (Den), Soma, Axon initial segment (AIS) and Distal Axon (DA)) based on the localization of Ankyrin G - an AIS specific scaffold protein (**Supplementary Fig. 8**). The trajectories were rotated to align along the ***x*** axis with anterograde transport pointing to the ‘+’ direction (**Supplementary Fig. 9, 10)**. Analysis of region-specific SVP/SV dynamics reveal several kinetic features that explain the steady-state SVP/SV distributions and target search mechanisms in neurons: *1)* we found that SVPs/SVs display compartment-specific search kinetics by dynamically partitioning between short-lived (30 ~ 80 ms) anterograde and retrograde transport movements interspersed by relatively long-lived slow diffusion states (100 ~ 200 ms; 0.06 ~ 0.09 μm^2^/s). (**Fig. 3b, c, e** and **Supplementary Fig. 11c**). *2)* In the AIS, the processivity of the anterograde transport is substantially enhanced, with anterograde transport lifetime and average movement velocity significantly increased (**Fig. 3c, d, e**). At the same time, the slow diffusion state has a substantially shortened lifetime in the AIS (**Supplementary Fig. 11c**). These observations suggest that specialized functional components in the AIS promote SVP anterograde transport while limiting transient binding and retrograde backtrack movements, in analogy to how a waterfall facilitates unidirectional water flows.*3)* Interestingly, in dendrites and distal axons, the strengths of anterograde and retrograde transport are balanced (**Fig. 3c, e** and **Supplementary Fig. 11a, b**), suggesting that SVPs unbiasedly sample these regions, dynamically searching for binding sites. The slow diffusion fraction is significantly higher in distal axons than in dendrites (**Supplementary Fig. 11c**), consistent with selective recruitment of SVPs/SVs to boutons and pre-synaptic regions. Previous studies show that the AIS consists of specifically arranged nanoscale cytoskeleton architectures^35, 36^ and the AIS serves as a passive filter controlling the selective entry of axonal cargos^37^. Data from our high-resolution kinetics analysis are complementary to these observations, highlighting a crucial role of the AIS in actively enhancing the axonal transport polarity by promoting the anterograde transport processivity (**Fig. 3e**).

### Imaging Long-lived Transcription Factor Binding Dynamics

We next sought to apply this method to imaging an entirely different molecular phenomenon - the binding of transcription factors (TFs) to chromatin. One highly debated area in this field is how long individual TF dwells on chromatin. Two popular single-molecule residence time measurement methods were developed recently: one is based on time lapse imaging^4, 38^ and the other relies on motion blur by using a long acquisition time of 500 ms^1, 3^. However, technical problems associated with dense labeling and photo-bleaching prevent these methods from reliably measuring the long-lived chromatin binding events at the single-molecule level. To address this problem, we devised an N-terminal RT stochastic labeling strategy, in which the RT sequence is placed upstream of the 3XHaloTag-TF sequence (**Fig. 4a**) and therefore protein over-expression is minimized.

**Figure 4:**
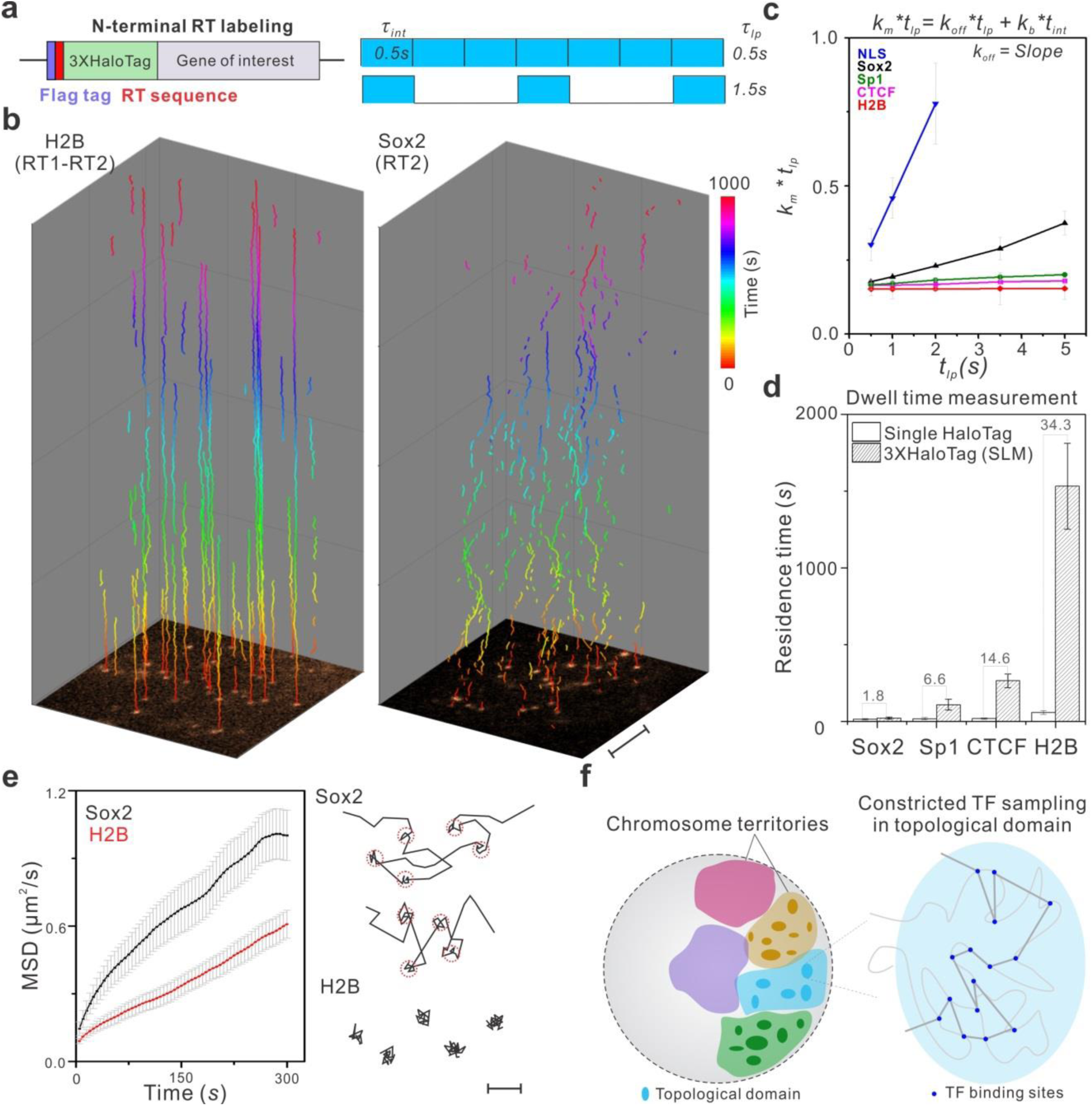
Long-time observation of TF-chromatin binding dynamics in the nucleus. (**a**) Left, the RT sequence is placed between a FLAG tag and a 3XHaloTag to achieve sparse labeling of transcription factors in live cells. Right, Due to the low labeling density, relatively long dark times between imaging frames can be introduced to probe long-lived binding events without tracking contaminations (See **Supplementary Fig. 12a** for details). Imaging integration time (*τ_int_*) is 500 ms; the lapse time 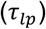 - the time between exposure starts can be varied. (**b**) With sparse labeling, single molecules are visible from the first frame shown at the bottom and can be faithfully tracked up to 1000 *sec* for H2B molecules (Left) with a lapse time of 5 *sec*. On the contrary, Sox2 binding is more dynamic (Right). In each trajectory, the time component displayed in the *z* direction is color-coded as indicated by the color bar. Imaging was performed using the Nikon TiE microscope (HILO illumination). Scale bar: 5 μm (**c**) By using a varied lapse-time imaging method as previously described^4^, the true chromatin dissociation rate (*k_off_*) of a TF can be calculated by using the slope of the linear regression of 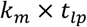 as a function of 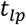. This calculation renders dissociation rate (*k_off_*) independent of photo-bleaching rate (*k_b_*). See **Supplementary Eqn. 1 - 8** for calculation details. *k_b_* is estimated as ~0.14 sec^−1^ according to **Supplementary Eqn. 8**. Tracking data from 8 ~ 10 cells were used in this analysis. Average number of trajectories for each data point: ~1850. H2B, SP1, CTCF and NLS imaging experiments were performed in U2OS cells. Sox2 experiments were performed in mouse ES cells. (**d**) Dwell time measurement comparison by using the traditional dense single HaloTag labeling strategy and the 3XHaloTag RT labeling strategy. The number above each bar reflects the fold of dwell time differences. (**e**) Left, the mean square displacement (MSD) plots show that, even when we selectively image less mobile fractions using long lapse times (5 *sec*) and a long acquisition time of 500 ms, Sox2 molecules are more mobile than H2B (Number of Trajectories: 1692 (H2B); 1049 (Sox2)). Error bars in (**c**), (**d**), and (**e**) represent SD. Right, a close examination of Sox2 and H2B trajectories shows distinct binding behaviors. Specially, Sox2 hops between adjacent stable binding sites (highlighted by 50 nm red circles). On contrast, H2B molecules remain relatively static at fixed locations. Scale bar: 100 nm (**f**) Long-term tracking suggests that Sox2 molecules translocate among binding sites in spatially-restricted regions inside the nucleus, supporting a sequestering mechanism of topological chromosome domains in regulating local gene activities.

After optimizing RT sequence for each TFs, we can visualize single molecules from the first imaging frame without high laser power pre-bleaching step (dSTORM) (**Fig. 4b** and **Supplementary Fig. 12b**). This is important for two reasons: *1)* because of the sparse labeling, in average only one labeled molecule exists in one diffraction limited area. Therefore, long dark times (0 ~ 4.5 *sec*) between imaging frames can be safely introduced without the risk of tracking errors where different nearby molecules across frames are linked as one (**Supplementary Fig. 12a**) *2)* our measurements do not rely on a single fluorophore switched back from the triplet state (dSTORM). More than one copies of JF549 dye molecules that are not pre-damaged ensure blinking/switching-proof, long-term observation of single-molecule dynamics (**Fig. 4b, Supplementary Fig. 12b, d, e** and **Supplementary Movie 8**). When we combined time lapse imaging (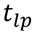, 0.5 ~ 5 *sec*) to minimize photo-bleaching and motion blur (500 ms exposure time) to selectively capture stable binding events (**Fig. 4a**), we were able to extend the temporal length scale of single-molecule dwell time measurement by ~2 orders of magnitude into the 10 ~ 20 mins range (**Fig. 4b, c, d** and **Supplementary Movie 8**). For example, average residence times for different classes of chromatin binding proteins (Sox2; 21.3 ±7.8 *sec*, Sp1; 109 ± 35.6 *sec*, CTCF; 265 ± 45.7 *sec* and H2B; 1532 ± 278.4 *sec*) measured by this approach are well separated in the temporal domain (**Fig. 4d**), consistent with their distinct roles in gene regulation. These results are in contrast to 12.1 ± 4.5 *sec* (Sox2), 16.5 ± 6.4 *sec* (Sp1), 18.1 ± 3.7 *sec* (CTCF) and 44.6 ± 12.6 *sec* (H2B) measured by the strategy described in previous reports^3, 39^.

One long-standing hypothesis in gene regulation is that topological chromosome domains might compartmentalize local gene activities by physically sequestering gene regulatory machinery. This notion was derived from extensive HiC (chromosome conformation capture coupled with high throughput sequencing) experiments^40^-^42^ and has yet to be substantiated by live cell data. Previously, it was shown that the stable binding sites of a pluripotency factor - Sox2 are spatially clustered in living embryonic stem (ES) cells^43^, suggesting the existence of high-order organization of cis-regulatory elements in the nucleus. However, due to technical problems associated with dense labeling and fast photo-bleaching, it is challenging to perform long-term tracking experiments to investigate whether Sox2 dynamics is altered in these binding site clusters. Here, long-time observation of slow diffusion events reveals distinctive dynamic differences between Sox2 and a chromatin marker (H2B) at the single-molecule level (**Fig. 4b**). Specifically, Mean Square Displacement (MSD) plots suggest that Sox2 molecules are more mobile than chromatin (H2B) even when we selectively imaged slow diffusion events by motion blur and long lapse times (2, 5 *sec*) (**Fig. 4e** and **Supplementary Fig. 13a**). A detail analysis of individual trajectories revealed that Sox2 dynamically hops and interacts locally in the nucleus (**Fig. 4b, e, Supplementary Fig.13** and **Supplementary Movie 8, 9**). In contrast, H2B molecules are generally more static, bound to fixed positions (**Fig. 4b, e** and **Supplementary Fig. 13a)**. We note that Sox2 molecules in the same field of view displayed both stable binding and hopping behaviors (**Supplementary Fig. 13b** and **Supplementary Movie 9**), excluding the possibility that the hopping is an artifact of cell movement or imaging platform drift. These results suggest that topological structures in the nucleus kinetically facilitate Sox2 molecules to explore locally, confining target search and gene regulatory activities (See **Supplementary Fig. 13a** and **Supplementary Eqn. 12, 13** for details of the confined motion analysis). Our studies thus provide the first evidence supporting a dynamic sequestering mechanism of topological structures in the nucleus of a live cell (**Fig. 4f**).

In summary, here we demonstrate a simple genetic handle to precisely titrate protein labeling density in live cells. Pairing with bright fluorescent labels, this method enables long-term single-particle tracking of densely-packed cellular organelles and proteins. We note that the strategy outlined here aims to unambiguously track one molecule as long-term as possible to probe its exquisite dynamic modes of action in live cells. Compared to sptPALM, this method trades off high trajectory densities for long trajectory lengths. However, if desirable, one could perform sptPALM while monitoring long-term dynamics in a two-color imaging experiment. The beauty of this method lies in its simplicity. Only conventional microscopy is required for long-term single-molecule imaging. However, this strategy can be readily combined with advanced imaging modalities^10^, such as lattice light-sheet microscopy^44^, adaptive optics microscopy^45^ and the MINFLUX method^11^ to study *in vivo* kinetics and control logics of molecular systems with extended imaging depth at higher spatiotemporal resolution.

## METHODS

### Cell Culture

Mouse D3 (ATCC) ES cells were maintained on 0.1% gelatin coated plates in the absence of feeder cells. The ES cell medium was prepared by supplementing knockout DMEM (Invitrogen) with 15% FBS, 1 mM glutamax, 0.1 mM nonessential amino acids, 1 mM sodium pyruvate, 0.1 mM 2-mercaptoethanol and 1000 units of LIF (Millipore). NIH/3T3 cells and U2OS cells were maintained in DMEM medium (Corning, without phenol-red) supplemented with 10% FBS, 1 mM glutamax. SH-SY5Y cells were cultured in DMEM/F12 medium (Gibco, without phenol-red) supplemented with 10% FBS.

Electroporation was used to transfect ES cells. Specifically, the Amaxa 4D-Nucleofector System and the kit for Mouse Embryonic Stem Cells (Lonza) were used according to manufacturer’s suggestions. We used Lipofectamine 3000 reagent (Invitrogen) for transfecting NIH/3T3 cells, U2OS cells and SH-SY5Y cells. For generating stable cell lines, cells were co-transfected with Piggybac transposon vector (neomycin-resistant) and a helper plasmid that encodes Piggybac transposase (Supper Piggybac Transposase, System Biosciences). Two days post-transfection, the cells were subjected to G418 (Thermo Fisher Scientific) selection.

### Primary Culture of Hippocampal Neurons

We prepared dissociated hippocampal neurons from P0 to 1 Sprague-Dawley rat pups. Briefly, the hippocampi were dissected out and digested with papain (Worthington Biochemical). After digestion, the tissues were gently triturated and filtered with the cell strainer. The cell density was counted. ~2.5 × 10^5^ cells was transfected with indicated constructs by using P3 Primary Cell 4D-Nucleofector X kit (Lonza). After transfection, neurons were plated onto poly-D-lysine (PDL, Sigma)-coated coverslips and maintained in NbActiv4 medium (BrainBits) at 37 °C for indicated days before imaging.

### Plasmids and Molecular Cloning

H2B, Sox2, and Sp1 cDNA were amplified from constructs used in our previous studies^3,47^. CTCF cDNA was amplified from ES cell cDNA libraries. Synaptophysin cDNA and 24xGCN4 (SunTag) cDNA were obtained from Addgene (Plasmid #51509 and Plasmid #72229). The scFv-GCN4-sfGFP-NLS, ankG-mCherry, and SEP-NR1 constructs were obtained from Addgene (Plasmid #60906, Plasmid #42566, and Plasmid #23999). The DNA fragment of 3XHaloTag with M to L substitutions was synthesized by Integrated DNA Technologies. The cDNA fragments were cloned into Piggybac transposon vector (PB533A-2, System Biosciences) or modified Piggybac transposon vector with human synapsin 1 promoter for neuronal specific expression. HaloTag (Promega) was used to replace the sfGFP region to get scFv-GCN4-halo-NLS construct. The plasmids used for fish microinjection were made using the Tol2-Gateway system and the pDEST vector was used as the destination construct. We note that the C-terminal RT strategy is best to combine with Cas9 knock-in to sparsely label protein from the endogenous locus. However, we can combine the N-terminal RT strategy with transgenic expression to achieve sparse labeling and at the same time minimize over-expression.

### Real time PCR

Total RNA was extracted with Trizol Reagent and reverse-transcribed by SuperScript III Reverse Transcriptase with oligo-dT primer (Thermo Fisher Scientific). cDNA corresponding to 5 ng of total RNA was used in each SYBR Select Master Mix (Thermo Fisher Scientific) reaction. Reactions were performed in duplicates and the results were collected on a CFX96 Touch Real Time PCR Detection System (Bio-Rad).

The PCR primers for H2B-mEOS4b-HA construct are:

Forward: 5-AATGTATGTGCGTGATGGAGTG-3

Reverse: 5-TATGGGTAACCTGAACCTGATC-3

The primers for human GAPDH are:

Forward: 5- AACGGATTTGGTCGTATTGGGC-3

Reverse: 5- CCTGGAAGATGGTGATGGGAT-3

### Western Blot

Whole cell extracts from U2OS cells were isolated using RIPA buffer that contained Complete Protease Inhibitor Cocktail (Roche). Protein concentrations were measured using Bio-Rad Protein Assay against BSA standards. Protein from each sample was resolved by SDS-PAGE. Primary antibodies used: HA tag (ab9134, Abcam, 1:500) and beta-tubulin (#2128, Cell Signaling Technology, 1:1000). HRP conjugated secondary antibodies (Pierce) were used at a dilution of 1:2000. LumiGLO chemiluminescent substrate (Cell Signaling Technology) was used for HRP detection and light emission was captured by films.

### Immunofluorescence Staining

Cells were first fixed with 4% paraformaldehyde, permeabilized and blocked with 10% goat serum, 1% BSA, and 0.25% Triton in PBS. Samples were stained with synaptophysin (ab8049, Abcam, 1:500) or synapsin 1 (AB1543P, Millipore, 1:5000) primary antibodies in PBS containing 10% goat serum, 1% BSA, and 0.1% Triton. Secondary antibodies: DyLight 488 conjugated secondary antibodies (Jackson ImmunoResearch, 1:400).

### Immuno-EM Experiment

Primary hippocampal neurons were grown on PDL-coated 8-well chamber slide (Thermo Fisher Scientific) for 10 ~ 14 days. Pre-embedding immunogold labeling was performed on the wells as previously reported^49^. Briefly, Neuronal cultures were fixed with 4% paraformaldehyde/0.05% glutaraldehyde in sodium cacodylate buffer (0.1 M, pH 7.2). After washing with cacodylate buffer and PBS, samples were blocked with 50 mM glycine in PBS and permeabilized in PBS containing 5% normal goat serum and 0.1% saponin. Then they were incubated with anti-GFP antibody (A-11122, Thermo Fisher Scientific, 1:200) first and Nanogold-Fab’ fragment of goat anti-rabbit IgG (Nanoprobes; 1:150) later. After silver enhancement (HQ kit, Nanoprobes) for 6 minutes, the samples were treated with 0.5% osmium tetroxide in phosphate buffer, dehydrated in ethanol and finally embedded in Eponate 12 resin (Ted Pella, Inc) for ultrathin sectioning. Ultrathin sections were contrasted with aqueous uranyl acetate and lead citrate and imaged on a Tecnai Spirit electron microscope (FEI, Hillsboro, OR) using an Ultrascan 4000 (Gatan, Inc) digital camera.

### Microinjection and imaging of *zebrafish*

Using a *zebrafish* gal4-vp16 driver line under the control of the pan-neuronal Huc promoter, embryos were co-injected at the 1 or 2-cell stage with plasmid DNA of UAS(5X)-syp-RT1-24xGCN4 at 25 ng/μl and 8 ng/μl for UAS (10X)-scFv-GCN4-sfGFP-NLS along with 25 ng/μl Tol2 mRNA. Fish were examined 48 ~ 60 hours later for GFP fluorescence and positive ones were picked to do the HILO imaging on a Nikon Eclipse TiE Motorized Inverted microscope equipped with a 100X Oil-immersion Objective lens (Nikon, N.A. = 1.49).

### Cell Labeling Strategy and Preparation for Imaging

Transfected hippocampal neurons were plated onto an ultra-clean cover glass precoated with PDL and cultured for indicated days (DIV 6 or DIV 12). For live imaging, culture medium was replaced with Tyrode’s solution (140 mM NaCl, 5 mM KCl, 3 mM CaCl_2_, 1 mM MgCl_2_, 10 mM HEPES, 10 mM glucose, pH 7.35).

Stable ES cell lines were plated onto an ultra-clean cover glass pre-coated with IMatrix-511 (Clontech). ES cell imaging experiments were performed in the ES cell imaging medium, which was prepared by supplementing FluoroBrite medium (Invitrogen) with 10% FBS, 1 mM glutamax, 0.1 mM nonessential amino acids, 1 mM sodium pyruvate, 10 mM HEPES (pH 7.2 ~ 7.5), 0.1 mM 2-mercaptoethanol and 1000 units of LIF (Millipore).

Transfected NIH3T3 and U2OS cells were plated onto an ultra-clean cover glass without coating. The NIH3T3 and U2OS imaging medium was prepared by supplementing FluoroBrite medium (Invitrogen) with 10% FBS, 1 mM glutamax, 0.1 mM nonessential amino acids and 1 mM sodium pyruvate.

For the saturated labeling of SunTag (24XHaloTag) or 3XHaloTag, cells were incubated with JF646-HTL or JF549-HTL with final concentration of 500~1000 nM for 30 mins. HTL stands for HaloTag Ligand. Chemical structures and synthesis procedures of JF549-HTL and JF646-HTL were described previously^5^.

To image single molecules without using the sparse labeling strategy, we first tested the optimal HaloTag-JF549 and HaloTag-JF647 labeling concentrations. Several concentrations of JF549-HTL and JF646-HTL (0.5 nM, 1 nM, 2 nM and 5 nM) were used to treat cells for 15 mins and then cells were washed with imaging medium for 3 times. The cover glasses were then transferred to live-cell culturing metal holders and mounted onto the microscope one by one. Proper HaloTag-JF549 or HaloTag-JF646 labeling concentrations were determined by the criterion that single-molecules can be easily detected after no or a minimal 2 ~ 5 *sec* pre-bleaching. After fixing the labeling concentration for each cell line, we then proceeded to perform the 2D single-molecule imaging experiments.

### PALM Imaging to Estimate RT Efficiency

Live-cell PALM imaging experiments were used to estimate the H2B-mEOS4b labeling densities (**Fig. 1b, c, Supplementary Fig. 3d** and **Supplementary Movie 1**). Specifically, imaging was performed on a Nikon Eclipse TiE Motorized Inverted microscope equipped with a 100X Oil-immersion Objective lens (Nikon, N.A. = 1.49), four laser lines (405/488/561/642), an automatic TIRF illuminator, a perfect focusing system, a tri-cam splitter, three EMCCDs (iXon Ultra 897, Andor) and Tokai Hit environmental control (humidity, 37 °C, 5% CO_2_). Proper emission filters (Semrock) were switched in front of the cameras for GFP, JF549 or JF646 emission and a band mirror (405/488/561/633; BrightLine quad-band bandpass filter, Semrock) was used to reflect the laser into the objective. For sparse single-molecule imaging, mEOS4b moiety was stochastically converted from ‘Green’ to ‘Red’ state with low-dose 405 nm illumination (5 ~ 10 W/cm^2^). Then, single molecules were imaged at 50 Hz using a 561 nm laser with the excitation intensity of ~1000 W/cm^2^. 5000 frames were collected for each cell. For each RT condition, data from 10 ~ 20 cells were collected.

### Imaging SVP Dynamics

SVP particles were imaged using TIRF with our Nikon Eclipse TiE set-up described above. For tracking SunTag-GFP labeled SVPs, we used a 488 nm laser with the excitation intensity of ~100 W/cm^2^. For tracking SunTag-HaloTag-JF549/646 labeled SVPs, we used a laser with the excitation intensity of ~50 W/cm^2^. For imaging HaloTag-JF549 labeled SVPs, we used a 561 nm laser with the excitation intensity of ~1 kW/cm^2^. An acquisition time of 20 ms was applied for all conditions.

### Single-molecule Imaging of Long-lived Chromatin Binding Events

For imaging immobile TF or H2B binding events in live nuclei, we used the same Nikon Eclipse TiE set-up described above. The TIRF illuminator was adjusted to deliver a highly inclined laser beam to the cover glass with the incident angle smaller than the critical angle. Thus, the laser beam is thus laminated to form a light-sheet (~3 μm) above the cover glass. The oblique illumination (HILO) has much less out-of-focus excitation, compared with the regular Epi-illumination. TFs labeled with 3XHaloTag-JF549 were imaged using a 561 nm laser with the excitation intensity of ~50 W/cm^2^. To specifically probe long-lived binding events, we used 500 ms imaging acquisition time to blend fluorescence signals from by fast diffusing molecules into the imaging background by motion blur^3, 39^. Because we only measure stable binding events at the focal plane by motion blur. Fast 3D motion should not affect the measurements. We also introduce long dark times (*τ_D_*) between imaging frames to minimize photobleaching as previously described^4^. The dark times vary from 0 *sec* to 4.5 *sec*. To minimize drift during imaging, we performed imaging in an ultra-clear room with the precise temperature control system. The environment control chamber for cell culturing was fully thermo-equilibrated and any devices that introduce mechanical vibrations are separated from the air table where the microscope body resides. We calibrated our imaging system with beads to confirm minimal drift during imaging (*xy* drift < 100 nm per hour).

### Single-Molecule Localization, Tracking and Diffusion Analysis

For 2D single-molecule localization and tracking, the spot localization (x,y) was obtained through 2D Gaussian fitting based on MTT algorithms^50^. The localization and tracking parameters in SPT experiments are listed in the **Supplementary Table 1**. The resulting tracks were inspected manually. The RT efficiencies (in %, **Fig. 1c** and **Supplementary Fig. 3d**) were calculated by normalizing single-cell localization density from each RT condition to the average density in the no RT control. Diffusion coefficients (**Fig. 4e** and **Supplementary Fig. 12c, 13a**) were calculated from tracks with at least 5 consecutive frames by the MSDanalyzer^51^ with a minimal fitting R^2^ of 0.8.

### HMM-Bayes Analysis of SVP Dynamics

The diffusion and transport states of individual SVP trajectories were analyzed by HMM-Bayes program^33^ with default parameters. Specifically, maximal 3 states can be inferred from one trajectory. There are possibilities for one or two states as well (*e.g.* **Fig. 2c, d**). In the HMM-Bayes analysis, the transport process is modeled as directed motion (V) with Brownian diffusion (D) according to the classical equation, <*r*^2^>= 4*D*Δ*t* + (*V*Δ*t*)^2^^46^. The program categorizes each step in the trajectory to either diffusion or transport processes and calculates diffusion coefficient as well as transport velocity. To separate anterograde and retrograde transport processes, we rotate the trajectory and align the neurite along the x axis with the anterograde transport point to ‘+’ direction. Specifically, for each trajectory, we manually defined one proximal and one distal point along the neurite and then the angle Θ between the vector (from the proximal to the distal point) and the ‘+’ direction of x axis was calculate by *atan2 ()* function in Matlab 2015a. Subsequently, the trajectory was rotated by - Θ. After the rotation correction, the transport processes bifurcated to anterograde and retrograde direction (**Supplementary Fig. 9, 10**). For each functional compartment (soma, dendrites, distal axon and axon initial segment) in the neuron, the average diffusion coefficients, transport velocities and life-time of different states (**Fig. 2d, 3c-e** and **Supplementary Fig. 11**) were calculated. In total, data from 20 ~ 30 cells were analyzed for each compartment. The standard deviation reflects cell-to-cell variations.

### TF Residence Time Analysis

To map stable bound sites in the slow acquisition (500 ms) and time lapse condition, 0.1 μm^2^/s was set as maximum diffusion coefficient (D_max_) for the tracking. The D_max_ works as a limit constraining the maximum distance (R_max_) between two frames for a particle random diffusing during reconnection. Only molecules localized within R_max_ for at least two consecutive frames will be considered as bound molecules. The duration of individual tracks (dwell time) was directly calculated based on the track length.

The relationship between photo-bleaching and dissociation can be described by the following diagram and differential equations, where *n*_0_ is the number of 3XHaloTag with 3 bright dye molecules at a given time; *n*_1_ is the number of 3XHaloTag with 2 bright JF dye molecules at a given time; *n*_2_ is the number of 3XHaloTag with 1 bright JF dye molecules at a given time; *k_off_* is the TF dissociation rate; *k_b_* is the photo-bleaching rate.

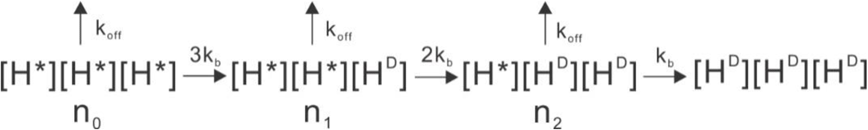

*H*^*^, HaloTag with a bright dye molecule. *H^D^*, HaloTag with a bleached dye molecule.

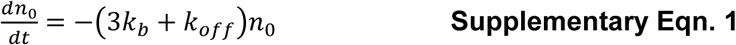

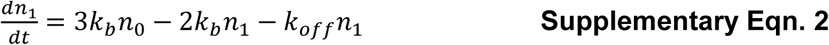

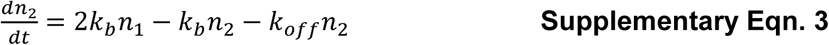

Here, we define *N_T_* as the number of 3XHaloTag labeled with JF dye molecules before imaging; *α* as the percentage (*N*_0_/*N_T_*) of triple-labeled molecules before imaging and *β* as the percentage (*N*_1_/*N_T_*) of double-labeled molecules before imaging. By solving the differential equations **1 - 3**, we can establish the relationship between the number of observable single molecules (*N_bright_* (*t*)), *k_off_*, *k_b_*, *α* and *β*.

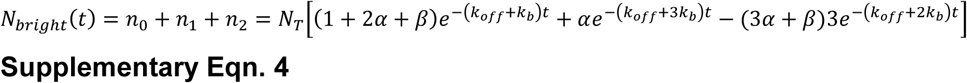

The time lapse photo-bleaching rate is equal to 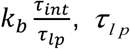 is the lapse time, *τ*_int_ is camera acquisition time. Thus **Supplementary Eqn. 4** can be transformed to

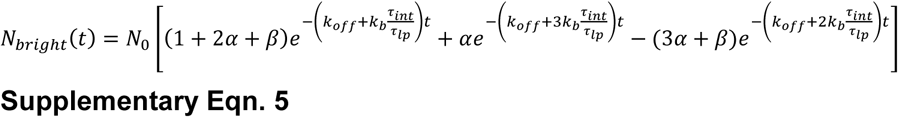

Here we further assign

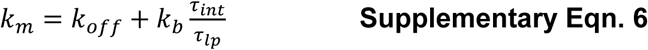

And, **Supplementary Eqn.5** can be rearranged to,

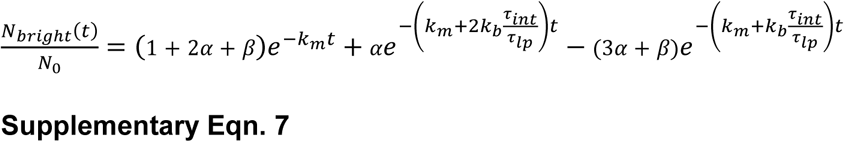

Fitting 1-CDF curve with **Supplementary Eqn.7** yields *k_m_*, *α* and *β* for different time lapse conditions. The fitting was performed using *lsqnonlin* function in Matlab 2015. **Supplementary Eqn.6** can be transformed into **Supplementary Eqn.8**

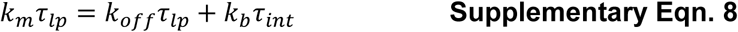

Therefore, the dissociation rate (*k_off_*) can be derived through linear fitting 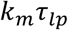 as a function of 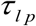 similar to the previous publication^4^.

### Jumping Angle Analysis

A sliding window of 3 points was applied to each track. The angle between the vectors of the first two and the last two points was calculated by the *acos()* function in the Matlab 2015a. Individual angles were pooled and binned accordingly for the angular Rose histogram (**Fig. 2d** and **Supplementary Fig. 10b**). The minimal jumping distance between two points is set as 40 nm to ensure that the angle measurement is not significantly affected by the localization uncertainty.

### Labeling Density Simulation and Motion-type Detection

For a 10 μm × 10 μm square area (**Supplementary Fig. 1a**) or for a 50 μm linear neurite (**Supplementary Fig. 5d**), the (x, y) positions for *N* single molecules were first randomly generated based on a uniform distribution in 2D or 1D. The photon counts contributed from each molecule to individual pixels (160 nm × 160 nm) were calculated based on the PSF estimator below.

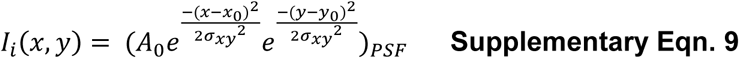

Specifically, *A*_0_ is the signal amplitude; σ is the Standard Deviation (S.D.: ~250 nm) of the Gaussian fit in the indicated direction, in our case S.D. of the x, y direction is the same.

To form the image, we first introduced random white noises (*B_xy_*) in the area with pixel intensities ranging from 0 ~ *A*_0_/5 (Signal-to-Background Ratio > 5). Then, the pixel intensity *I*(*x, y*) was calculated by iteratively summing the photon counts contributed from each molecules above the background value, according to equation below:

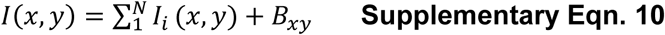

In **Supplementary Fig. 1**, we used the Einstein’s Brownian diffusion equation to calculate the expected jumping distance between frames.

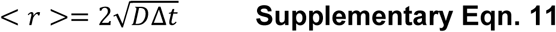

Note that when the motion is confined, the MSD curve is situated below its tangent at Δ*t* = 0. Therefore, if we model coarsely the MSD curve by a power law according to 2D diffusion models^46^.

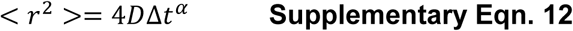

We should get α = 1 for purely diffusive motion, and α < 1 for confined motion. So we could determine from our experimental data a power law coefficient. This is best made in a log-log fashion (**Supplementary Fig. 13a**), for which power laws turn to linear laws:

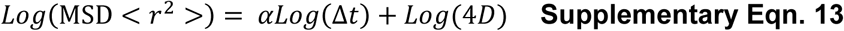

### Numerical Simulation of Protein Copy Number in the Cell

Because the number of reacting DNA molecules is small, random molecular fluctuations become significant and discrete, Gillespie stochastic simulation algorithm (SSA) approach^52^ is applied in this case. Such a system can be modeled as a continuous-time Markov process, whose probability distribution obeys what is called a chemical “master equation”. We used the Gillespie’s Direct Method for the simulation. We first provided a model consisting of a matrix of reaction rates, a function that computes reaction propensities (probability/unit time), an initial state vector, and initial/final times. The SSA functions return the sequence of event times and protein/RNA species amounts.

The master gene expression model is defined by the following reaction and parameters:

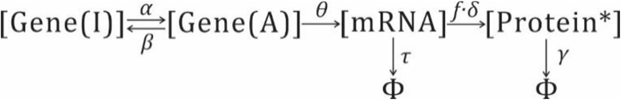

*α*, Gene Activation Rate
*β*, Gene Inactivation Rate
*θ*, mRNA Production Rate
*δ*, Translation Rate
*f*, Read-Through Rate
*δ*, Translation Rate
*α*, mRNA degradation Rate
*γ*, Protein degradation Rates
*, Labeled Protein

According to previous modeling calculation^53^, the mean copy number of labeled proteins is given by:

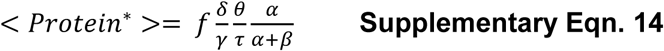

In the simulation, we kept mRNA Production Rate (*θ*), mRNA degradation Rate (*α*), Translation Rate (*δ*) and Protein degradation Rate (*γ*) constant. We used a RT Rate (*f*) of 0.5%, similar to the efficiency of RT3. We adjusted the promoter strength by tuning the fraction of the promoter on time (%), which is equal to *α*/(*α* + *β*). For reliable single-molecule tracking, we maintained the average protein copy number in the simulation as ~40 copies per cell. Here we defined the protein noise (*η*) as the standard deviation divided by the mean.

### Statistics

Comparisons between two groups were performed with Student’s *t*-test. Comparisons among multiple groups were performed with one-way ANOVA (**Supplementary** Fig.11c). Error bars in all figures represent SD, except the box charts. Differences were considered to reach statistical significance when *p* < 0.05.

### Data Availability Statement

The data that support the findings of this study are available from the corresponding author upon request.

## ACKNOWLEDGEMENT

We thank M. Radcliff, S. Moorehead and C. Morkunas for assistance. We thank M.G. Paez-Segala in Looger lab at Janelia research campus for mEOS4b construct. We thank Y. Liang, R. Turcotte in Ji lab and R. Chhetri in Keller lab at Janelia research campus for helping us test imaging conditions. This work is solely funded by Howard Hughes Medical Institute (HHMI).

## AUTHOR CONTRIBUTIONS

Z.L. and H.L. conceived and designed the experiments. H.L. performed and participated in all experiments. P.D., J.S. and M.K. performed the *zebrafish* experiment. P.D., M.S.I. and L.L. participated in the material preparation and initial test. H.A.P. and P.R. performed the EM experiments. J.G. and L.D.L. performed the JF dye synthesis. Z.L. and H.L. analyzed the data and wrote the paper. Z.L. supervised the research.

